# Effects of ploidy and genetic diversity on competitive outcomes

**DOI:** 10.1101/2023.02.23.529645

**Authors:** Jesús Alberto Pérez-Romero, Ana García Muñoz, Enrica Olivieri, A. Jesús Muñoz-Pajares, Mohamed Abdelaziz, Oscar Godoy

## Abstract

- Genetic diversity affects evolutionary trajectories but their ultimate effects on ecological interactions and community dynamics remains poorly understood. It has been hypothesized that phenotypic novelties produced by ploidy and heterozygosity modify the ecological interactions between novel genotypes and more ancient locally adapted ones, and therefore, their opportunities to coexist.
- We performed a greenhouse competition experiment with three taxa of the *Erysimum incanum* species complex differing in ploidy (2x, 4x and 6x) and heterozygosity (high and low). This experiment allows us to parameterize a population model to test the effect of genetic diversity on modulating the ecological forces that determine the outcome of competition, niche and fitness differences.
- Depending on whether ploidy variation and the level of heterozygosity made interspecific competition greater or smaller than intraspecific competition, we predicted either priority effects or coexistence. Such competitive outcome differences were explained by the phenotypic expression in the number of stalks (plant size surrogate) with genotypes under priority effects showing more stalks.
- Altogether, our results show that non-polyploid plants can coexist with polyploids contravening theoretical expectations of polyploidy dominance under stable conditions. However, historical contingency such as order of arrival promotes priority effects when adaptive phenotypic optimums strongly compete for space.

## Introduction

Progress in ecological theory during the last decades has substantially rendered a mechanistic understanding of the rules governing the maintenance of species diversity. These advances, formally named as modern coexistence theory (MCT) (Chesson, 2000), posits that there are two species differences that determine the outcome of competitive interactions. On the one hand, niche differences occurring when intraspecific competition exceeds interspecific competition stabilizes the population dynamics of interacting species by limiting their population growth when they are abundant but buffering them from extinction when they are rare (Adler *et al*., 2007). These stabilizing niche differences can arise from a wide variety of ecological factors such as differences in phenology (Godoy & Levine, 2014), differences in natural enemies (Petry *et al*., 2018), or differences in nutrient requirements (Harpole *et al*., 2016). On the other hand, fitness differences drive competition dominance and in the absence of niche differences determine the competitive superior. Fitness differences, understood within an ecological context, occur when good light competitors grow at the expense of other species (DeMalach *et al*., 2017) or when species are able to draw down common resources faster than the competitors (Tilman & Sandhu, 1998), and it is the result of two components. The first component is the species demographic differences, which arise from different ability of species to produce viable offspring, and the second component is the competitive response differences, which arises when species show different responses to competition. At the extremes, a species can be a superior competitor either because it produces a great amount of viable offspring or has a low sensitivity to competition (i.e. offspring production is not reduced when density increases), although a combination of both strategies is also possible.

Importantly, theory predicts a variety of outcomes depending on the relationship between niche and fitness differences (Ke & Letten, 2018). Under negative density-dependence (i.e., population growth rates decrease as the density of a population increases), species are predicted to coexist when niche differences overcome fitness differences. On the other hand, if fitness differences are overwhelming, the inferior competitor species are predicted to be excluded. It can also be the case that species are experiencing positive density-dependence (i.e., population growth rates increase as the density of a population also increases). In such cases, priority effects are expected to occur. This means that contingency processes such as order of arrival influence community assembly and the species that arrive first dominates the community and excludes the other (Fragata *et al*., 2022).

Modern coexistence theory was developed within an ecological context and as such most of its application has been done within this domain. This implies that the role of evolution in determining the outcome of ecological interaction is still poorly understood. Empirical work at the macroevolutionary scale has shown that disparate evolutionary processes among species poorly predict the outcome of ecological interactions and they can either determine coexistence or competitive exclusion (Narwani *et al*., 2013; Godoy *et al*., 2014; Germain *et al*., 2016). Moreover, Germain *et al. (*2016) showed that the scaling of niche and fitness differences with phylogenetic relatedness depend on whether species have evolved in sympatry or allopatry, being allopatric species less likely to coexist based on phylogenetic distance. At the microevolutionary scale, some examples have documented that rapid evolution ameliorates the negative effect of competition and ultimately can favor the coexistence of competing species (Lankau *et al*., 2009; Hart *et al*., 2019), whereas others have documented the opposite result (Qin *et al*., 2013). This lack of knowledge and context dependency calls for further studies to better mechanistically understand the effect of evolution on ecological interactions. In that regard, processes affecting genetic diversity have been long thought to be an important driver of ecological interactions, and it has been amply discussed that common processes should control the maintenance of both genetic and species diversity (Dempster, 1955; Ayala & Campbell, 1974; Hughes *et al*., 2008). However, detailed experiments to these this hypothesis are still lacking.

Two evolutionary processes are expected to modulate the degree of genetic diversity. The first and most important one is polyploidization, which is present in nearly 70% of flowering plants (Wood *et al*., 2009) and is playing an essential role in their evolutionary history (Grant, 1981; Soltis & Soltis, 1999; Soltis *et al*., 2009) and their diversification (Leebens-Mack et al. 2019). Complete genome duplication stimulates the neofunctionalization of duplicated, redundant genes, potentially leading to novel and innovative traits promoted by natural selection (Otto & Whitton, 2000; Parisod *et al*., 2010). Furthermore, it is broadly documented that genome duplications lead to variation in plant phenotype (Jürgens *et al*., 2002), involving changes in the rest of ecological interactions within a community. For example, an increment in flower size exhibited by higher ploidies might modify pollinator preferences between co-occurring individuals differing in ploidy level (te Beest *et al*., 2012; Moghe & Shiu, 2014). Ultimately, polyploidization events are able to change the resources usage and, thus, how diploids and their polyploid counterparts are spatially located (Levin, 1981; Raabová *et al*., 2008; Kolář *et al*., 2013). The overall increasing fitness in polyploid species is suggested as a potential driver for ecological adaptation to colonize novel habitats and face a major diversity of environmental conditions compared to diploids, which would explain the diversification patterns shaped by polyploids, especially in islands (Meudt *et al*., 2021). In sum, these previous findings suggest that ploidy is a driver of both fitness and niche differences but empirical assessments that explicitly explore how these differences determine the outcome of competition are lacking.

Together with polyploidization, a second important characteristic is the degree of heterozygosity. Heterozygosity is key in understanding the ecological consequences of competing genotypes because, just like polyploidy, it also increases raw material in the long term by novel allele combinations for evolution to act upon (Nieto Feliner *et al*., 2020). The allelic diversity effect is shown through changes in phenotype and even in the individual performance. An example is the classical heterosis event exhibited by the offspring originated by outbreeding (Hayes & Others, 1952; Bomblies & Weigel, 2007). This occurs when the heterozygotic offspring resulting from outbreed crosses exhibit a major performance compared with homozygotic parents. However, heterosis has been well documented in crop plants because heterozygotic phenotypes are commonly accompanied by a higher performance and adaptive ability (Fridman, 2015). Studies in heterosis help to understand the genotype-phenotype relationship due to the presence of different alleles resulting in phenotypes that, ultimately, could be able to drive evolutionary processes. However, both polyploidization and heterozygosity has been also shown to produce an immediate effect on the individual fitness within a single generation (Ramsey & Schemske, 2002). For this reason, comparisons of fitness differences between homozygotes and heterozygotes or diploids and polyploids, are commonly investigated to explain their coexistence or spatial segregation (Sonnleitner *et al*., 2010; te Beest *et al*., 2012; Ramsey & Ramsey, 2014).

If we summarize all these previous findings on the effect of genetic diversity on ecological dynamics, we can posit that they have been mostly focused on understanding what processes drive the fitness differences among genotypes that determine competitive exclusion. (Ramsey & Schemske, 2002). Within this perspective, coexistence has been considered a spatial process in which different genotypes persist under different locations thanks to being locally adapted. However, MCT predicts that the persistence of genetic diversity can be also achieved within the same location by promoting niche differences that stabilize the dynamics of competing genotypes. Yet, information on how genetic diversity promotes these niche differences is currently missing (Rey *et al*., 2017). Including the axis of niche differences is critical to understand when new variants are able to coexist with their ancestors within the same location when genome duplication occurs, or when the new variant excludes (or it is excluded by) the ancestor. This is well illustrated in the case of many species as the case of strawberry, which have evolved their genome in response to arid or stressful conditions (Liston *et al*., 2020). We can hypothesize that if genome duplication has served to cope with stressful conditions, then, there is likely to observe niche differences among genotypes with different ploidy due to niche segregation. Likewise, genetic diversity can also promote niche differences by phenotypic changes that allow new variants to explore different resources(Kolář *et al*., 2013); (Hernández-Leal *et al*., 2019). Overall, we have expectations that genetic diversity promotes both niche and fitness differences and the study of the effect of genetic diversity on the drivers of competitive outcomes can allow us to obtain a better understanding of how microevolutionary processes maintain genetic diversity, or if this does not occur, it allows to identify which genotypes are excluded by deterministic processes because they become inferior competitors after evolutionary processes or by contingency due to the order or arrival (i.e., priority effects)(Fig. 1). Fortunately, there are tools readily available to explore these questions by combining population models with detailed experiments in which it is possible to measure fitness and density dependent processes (Narwani *et al*., 2013; Godoy & Levine, 2014; Germain *et al*., 2016).

**Figure 1.**
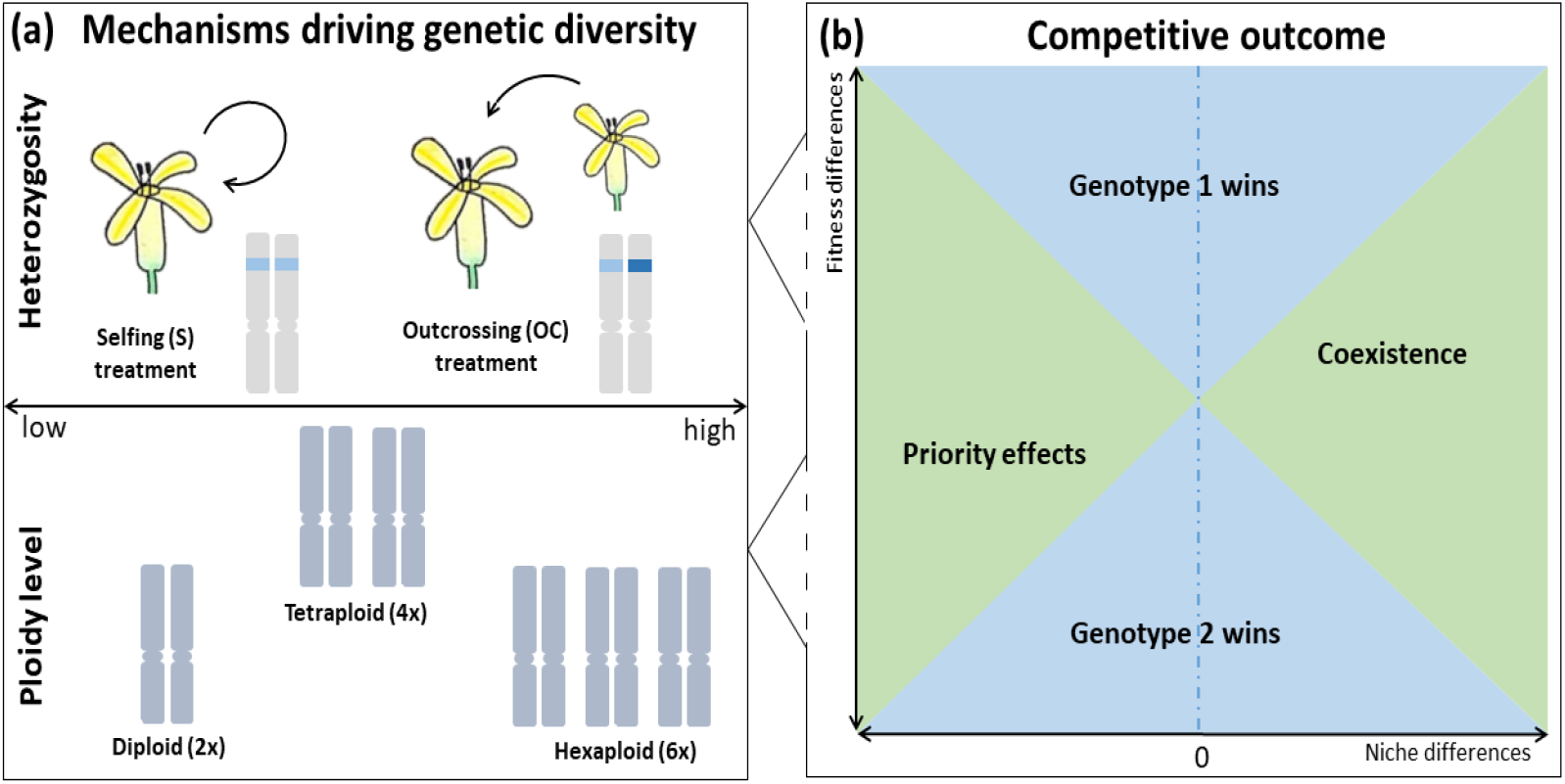
(a), Mechanisms driving the genetic diversity of the studied individuals, and (b) competitive outcomes expected in our experiments according to niche and fitness differences. The genetic diversity of the study system is mainly affected by the ploidy level (including diploid, tetraploid and hexaploid individuals) and by the heterozygosity level (the studied individuals have been produced after performing selfing and outcrossing manual pollination on parental individuals). Genetic diversity is expected to increase with ploidy level and outcrossing treatments. Such genetic diversity is expected to influence in turn the outcome of competition, depending on how they promote niche and fitness differences. Three different outcomes are expected. Coexistence (green area) in which both genotypes do not exclude each other, competitive exclusion (orange area) in which the genotype with higher fitness excludes the other, and finally, priority effects (blue area) in which the genotype that arrives first exclude the other.

Here, we focus on the annual multiploidy species complex *Erysimum incanum* (Brassicaceae) to address how genetic diversity determined by the degree of polyploidization and heterozygosity influences ecological interactions and competitive outcomes between contrasted genotypes. This variation in genetic diversity was obtained by combining different genotypes from diploids to tetraploids and hexaploids with crosses among individuals within the same level of ploidy to increment the degree of heterozygosity (Fig. 1). With this experimental set up, we were able to answer the following questions: 1) Does ploidy determine differences in fitness among genotypes? 2) How does the interaction between ploidy and heterozygosity influence the niche and fitness differences that determine the outcome of competition between genotypes and the maintenance of genetic diversity?, 3) Is a particular character able to summarize variation in competitive outcomes among genotypes?

## Material and Methods

### Study system and experimental set-up

We focus our study in the genus *Erysimum L*., which is one of the most diverse in the Brassicaceae family, with species inhabiting Eurasia, North Africa and North and Central America (Al-Shehbaz *et al*., 2006). In particular, we studied the species complex *Erysimum incanum*. This complex includes annual monocarpic species and subspecies inhabiting the Western Mediterranean basin, which is a main diversification center of the genus (Abdelaziz *et al*., 2011; Nieto Feliner, 2014). Within this complex and using flow cytometry, we found three ploidy levels: diploids (2x = 16 chromosomes), tetraploids (4x = 32) and hexaploids (6x = 48) (Nieto Feliner, 2014; García-Muñoz *et al*., 2022) with dissimilar geographic distribution. Diploids of *E. incanum* (*Erysimum incanum subsp. mairei*), present a vicariant distribution between the Rif and the Pyrenees mountains while tetraploids (*E. incanum subsp. incanum*) present a similar distribution in southwest Iberian Peninsula and the Middle Atlas Mountains (Fennane & Ibn-Tattou, 1999). In contrast to these ploidies occurring in both continents, hexaploid plants (*Erysimum meridionalis sp. nov*.) has been only found in the High Atlas and Antiatlas mountains (Abdelaziz et al., *in prep*.). Most species of the *E. incanum* complex exhibit autogamy as the predominant reproductive strategy, showing hermaphroditic flowers with the specific characteristics of the selfing syndrome. This reproductive system results in full-sib individuals within the same family.

Using this species complex as a baseline, we removed any local effects before performing experiments by obtaining pure lines from more than five generations in controlled conditions. Once these pure lines were obtained, we further modified their degree of genetic diversity within each ploidy level by crossing individuals in order to maintain or remove the homozygosity exhibited by pure lines. To do so, the first treatment consisted in selfing hand-made crosses that allow obtaining seeds in which the homozygosity level is assumed to be complete within the pure line (S). The second treatment consisted in intra-population allogamous crosses, where some flowers of each plant were emasculated before first opening and pollinated with pollen of individuals from a different family within the same population. This latter treatment resulted in seeds in which homozygosity was replaced by a full degree of heterozygosity (OC), except for these loci where alleles were identical by state. Overall, this procedure led to three “selfing (S)” and three “outcrossing (OC)” plant families according to the low or high degree of heterozygosity, respectively, which were factorially combined with the three ploidy levels. Therefore, a total of 18 plant families, three per combination of 2xS, 2xOC, 4xS, 4xOC and 6xS, 6xOC were used for evaluating experimentally the role of genetic diversity in coexistence outcomes.

### Theoretical approach

Our greenhouse experiments were designed to experimentally parameterize a mathematical model describing the population dynamics of interacting species (Levine & HilleRisLambers, 2009), which here was extended to genotypes. With this model, it is possible to quantify stabilizing niche differences and average fitness differences between interacting organisms from plants to animals (Godoy & Levine, 2014; Fragata *et al*., 2022). The model is described as follows:

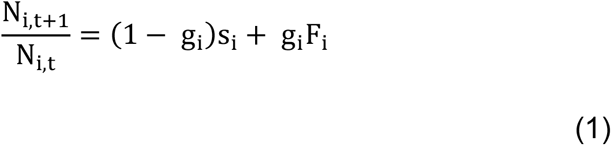

Where 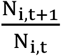 is the per capita population rate, and N_i,t_ is the number of seeds of genotype *i* in the soil prior to germination in winter of year *t*. In addition, the germination rate of species *i*, s_i_, can be viewed as a weighting term for an average of two different growth rates: the annual survival of non-germinated seed in the soil (g_i_), and the viable seeds produced per germinated individual (F_i_). We assume that genotypes affect the performance of one another when germinated individuals limit the fecundity of competitors. Thus, the per-germinant fecundity, F_i_, can be expanded into a function including the density of competing individuals in the system.

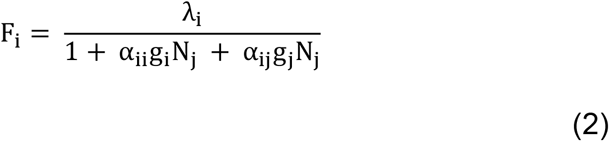

where λ_i_ is the per-germinant fecundity in the absence of competition. It is reduced by germinated individuals of its own and other species, which are multiplied by interaction coefficients, α_ij_, that describes the per capita effect of genotype j on genotype i. The model ignores the potential for age-dependent survival of non-germinated seeds, because prior work in annual plants has shown that seed bank survival has negligible influence on the competitive outcomes (Godoy & Levine, 2014).

With the dynamics of competition among genotypes described by this population model, we followed the approach of (Chesson, 2012)) to determine fitness and niche differences between species pairs. Following (Godoy & Levine, 2014) method, niche overlap between pair of genotypes, ρ, was calculated as:

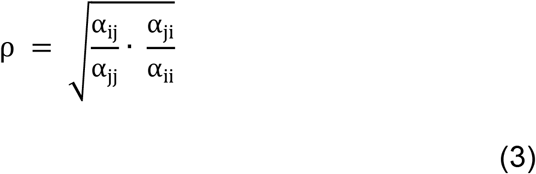

Niche overlap describes the degree to which competition among individuals of the same genotype (α_ii_, α_jj_) impact more than competition among individuals of different genotypes (α_ij_, α_ji_). Niche overlap span from zero (i.e., no niche overlap) to one (i.e., complete niche overlap). With (ρ) defining niche overlap between a pair of genotypes, their stabilizing niche difference is expressed as 1−ρ. As hypothesized, we expect that genetic differences in ploidy and in heterozygosity and their combination will reduce niche overlap among genotypes, and therefore, will increase niche differences.

As an opposing force to stabilizing niche differences, average fitness differences drive competitive dominance, and in the absence of niche differences, determine the competitive superiority between a pair of genotypes. Following previous methodologies (Godoy & Levine, 2014), we define average fitness differences between the competitors (ji) as:

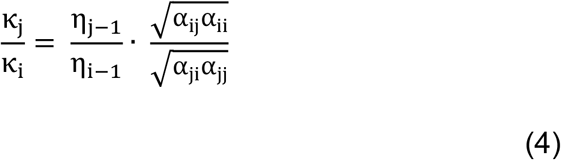

where η_i_ and η_j_ are the annual seed production for both genotypes and α_ji_ and α_jj_ are the per capita effect of a genotype i and genotype j on the seed production of a genotype j, respectively. It is worth noting that we did not explicitly estimate the germination rates (g_i_) and soil survival rates (s_i_) but we consider them to be equal to one. Therefore, in this particular study η_i_ and η_j_ are equal to λ_i_ and λ_j_ respectively. According to equation 4, average fitness differences can be decomposed in two different expressions. On the one hand 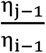 describes the “demographic difference” (i.e., the extent to which genotype i produces more seeds per germinant than genotype j). On the other hand, 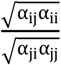 describes the “competitive response ratio” (i.e, the extent to which genotype i is more sensitive to competition than genotype j). From the expression of average fitness difference (equation 4), we can describe the genotype competitive ability (Hart *et al*., 2018)

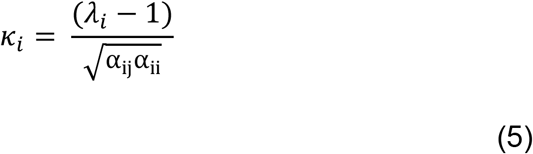

The competitive ability (*k*_*i*_) describes the ability of a genotype to be a superior competition as a function of two possibilities. Either because it can produce a high amount of viable seeds (*λ*_*i*_ − 1) or the genotype is not sensitive to competition with other genotypes 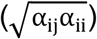, that is the amount of viable seeds produced is not reduced as the density of the competitor increases.

Importantly, the greater the ratio between genotypes j and i, the greater the fitness advantage of genotype j over i. If this ratio is one, genotypes are equivalent competitors. Coexistence requires both genotypes to invade when rare (Chesson, 2012). Then we established coexistence condition as (Godoy & Levine, 2014):

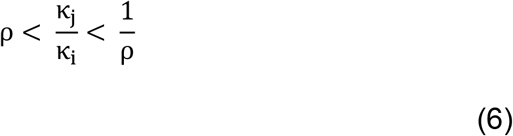

This condition allows us to distinguish three coexistence outcomes. The first outcome is *stable coexistence* that occurs when niche differences are larger than fitness differences. The second outcome is *competitive exclusion* that occurs when fitness differences are larger than niche differences. And finally, the third outcome is *priority effects* that occur when niche differences are negative, which indicates that species are experiencing positive density-dependence. In that final outcome it is predicted that the species that arrives first to the community excludes its competitor (Ke & Letten, 2018) (Fig. 1).

### Competition experiment and character measurements

To empirically parameterize the population model with which we can determine the competitive outcomes between genotypes, we conducted greenhouse experiments to estimate per germinant fecundities in the absence of neighbors (λ_i_), and all pairwise interaction coefficients (α_ij_). In March 2020, we displayed pots of 4.2 L (0.18×0.18×0.13cm), which were filled with Gramoflor™ potting soil mixture and watered every two days. The overall design involved sowing each genotype as focal individuals into a density gradient of each competitor genotype (including itself). To create this gradient, we followed a spatial explicit design within each pot proposed by (Bartomeus *et al*., 2021), in which focal genotypes of the same family experienced a density gradient from 1 to 4 individuals of a different genotype family. In order to calculate all pairwise interaction coefficients (α_ij_), this density gradient was created for each pairwise combination of families. We also grew individuals alone to better estimate the fecundities in the absence of neighbors (λ_i_). To estimate such fecundity that we understand in an ecological context is the “demography performance” of the genotype and in the evolutionary context is the “fitness” of an individual, we counted the number of fruits per plant and multiplied it by the mean number of viable seeds estimated from four random fruits in the same plant. This way, we obtained the total viable seed production per individual plant. This value is an unbiased estimate of the individual fitness due to the monocarpic life form of the study system.

Together with the competition experiment, we measured a series of characters for each family at the peak of individual biomass. These characters were related to the vegetative body plant overall related to vegetative body plants. Specifically, for every focal individual that produced seeds we measured its plant height, the number of flowering stalks and the diameter of the main stalk. This was done for a total of 210 individuals with 35 individuals per family.

### Statistical analyses

We used maximum likelihood techniques to parameterize the population model following a nested approach. That is, we first created a single model for which we estimate the intrinsic growth rate in absence of competitors (λ_i_), and then we used this information as prior for subsequent more complex models that include an overall term of competition in the second step and intra and interspecific competitive interactions (the *α*’s) in the third final step (Matías et al., 2018). λ_i_ were considered fixed per genotype family species but competition varied across genotype pairs. Finally, we used a one-way ANOVA in order to test whether coexistence outcomes between genotype pairs could be explained by a particular plant character. All analyses were done using R (R Core Team, 2021). To predict coexistence outcomes, we used the package ‘cxr’ (García‐ Callejas *et al*., 2020). Plots were done using ‘ggplot2’ (Wickham, 2016) and ‘cowplot’ (Wilke, 2019) packages.

## Results

Our results show that the genetic diversity of the different genotypes contribute to promote differences in viable seed production as well as competitive interactions. When we decomposed average fitness differences into its demographic and competitive response components, we found that diploids (2x) were the most competitive genotypes followed by tetraploids (4x), and finally hexaploids (6x) under the experimental conditions we imposed with no drought treatment (Fig. 2A). This competitive superiority of the diploids was due to a higher viable seed production in the absence of competition as well as lower sensitivity to reduce viable seed production in the presence of neighbors (Fig. 2B and 2C). Conversely, the low competitive ability of the hexaploids were due to a combination of lower viable seed production and higher sensitivity to competition. The amount of niche differences between the diploids and the other ploidy levels was not enough to overcome their observed differences in average fitness. These results overall indicate that diploid genomes are the superior competitor. However, they were not able to competitively exclude the other ploidy families (Fig. 3A). This outcome is driven by the low differences in fitness and response to competition (Fig. 2), which lead diploids to share the scenario of strong priority effects with the two other ploidies

**Figure 2.**
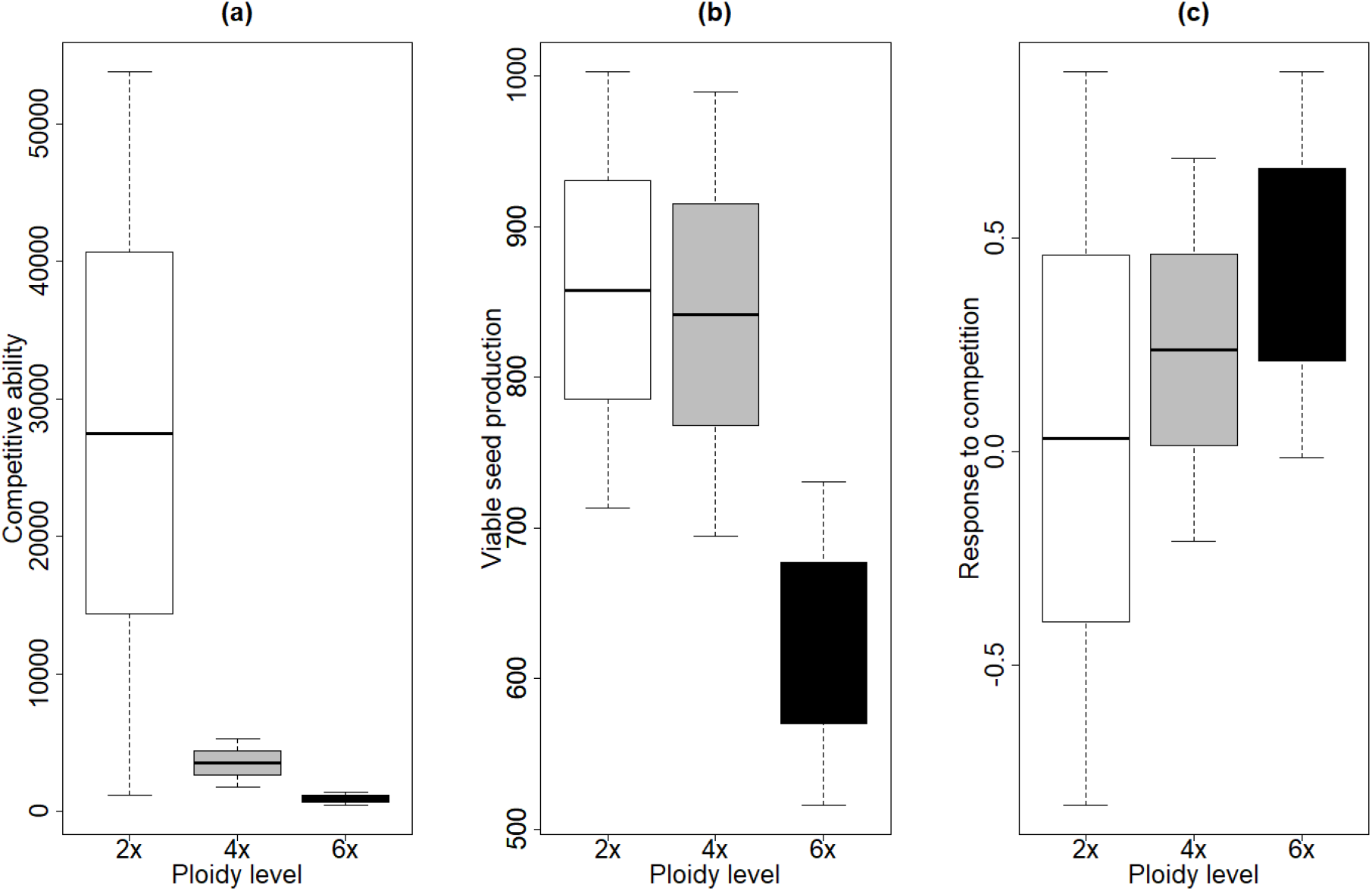
Competitive ability (A), viable seed production (B) and response to competition (C) for each one of the three ploidy levels tested in *E. incanum* system.

**Figure 3.**
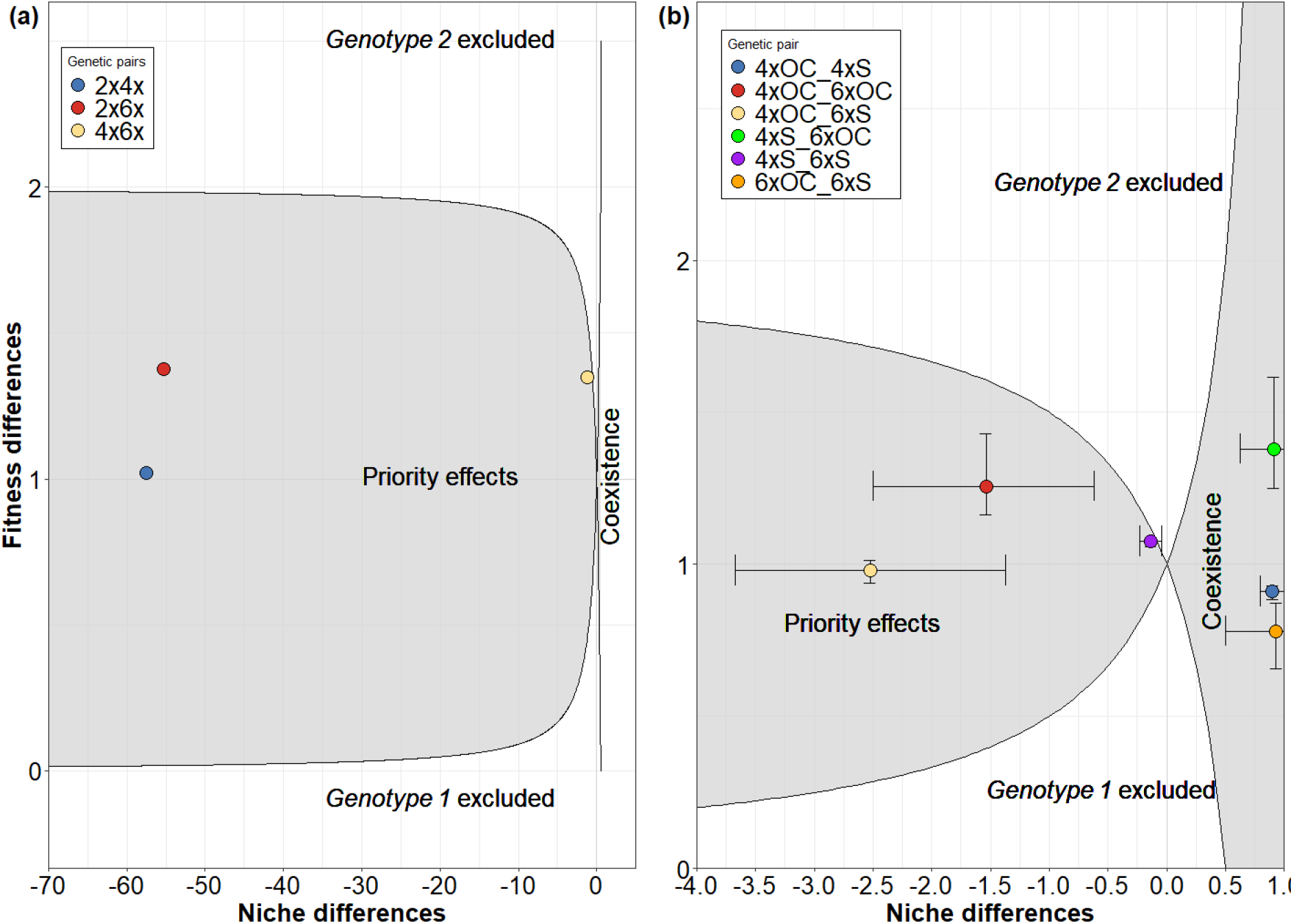
Relationship between fitness difference and niche difference for different combinations of ploidy (2x, 4x and 6x) in *E. incanum* system (A) and relationship between fitness difference and niche difference for different combinations of ploidy (4x and 6x) and levels of heterozygosity (low, S and high, OC) in *E. incanum* system (B). The two solid black lines represent the coexistence condition and its symmetrical for each ploidy level tested and defined the space in which genotypes could coexist and in which there were priority effects. Error bars show coexistence outcomes at the 95% confidence interval.

Such strong priority effects were not predicted when considering tetraploid and hexaploid genotypes in combination with their degree of heterozygosity. For these two levels of ploidy, we found two contrasted clusters of outcomes when we added more resolution by explicitly accounting for the degree of heterozygosity. One cluster in which priority effects among the pairs of genotypes were predicted to occur, and another cluster in which coexistence was predicted between three other different pairs of genotypes (Fig. 3B). This result was not predicted by theoretical expectations and suggests that genetic diversity produces a wider variation of ecological outcomes than previously expected.

Priority effects occur under positive density dependence when interspecific competition is stronger than intraspecific competition, whereas, coexistence occurs under negative density dependence when intraspecific competition is stronger than interspecific competition. Therefore, the change from one location of the coexistence map to another can be due to a change in intraspecific competition, interspecific competition or a combination of both. In our experiment, detailed analysis revealed that changes in the strength of interspecific interactions rather than intraspecific interactions were the main driver of switches from priority effects to coexistence regions (Fig. 4).

**Figure 4.**
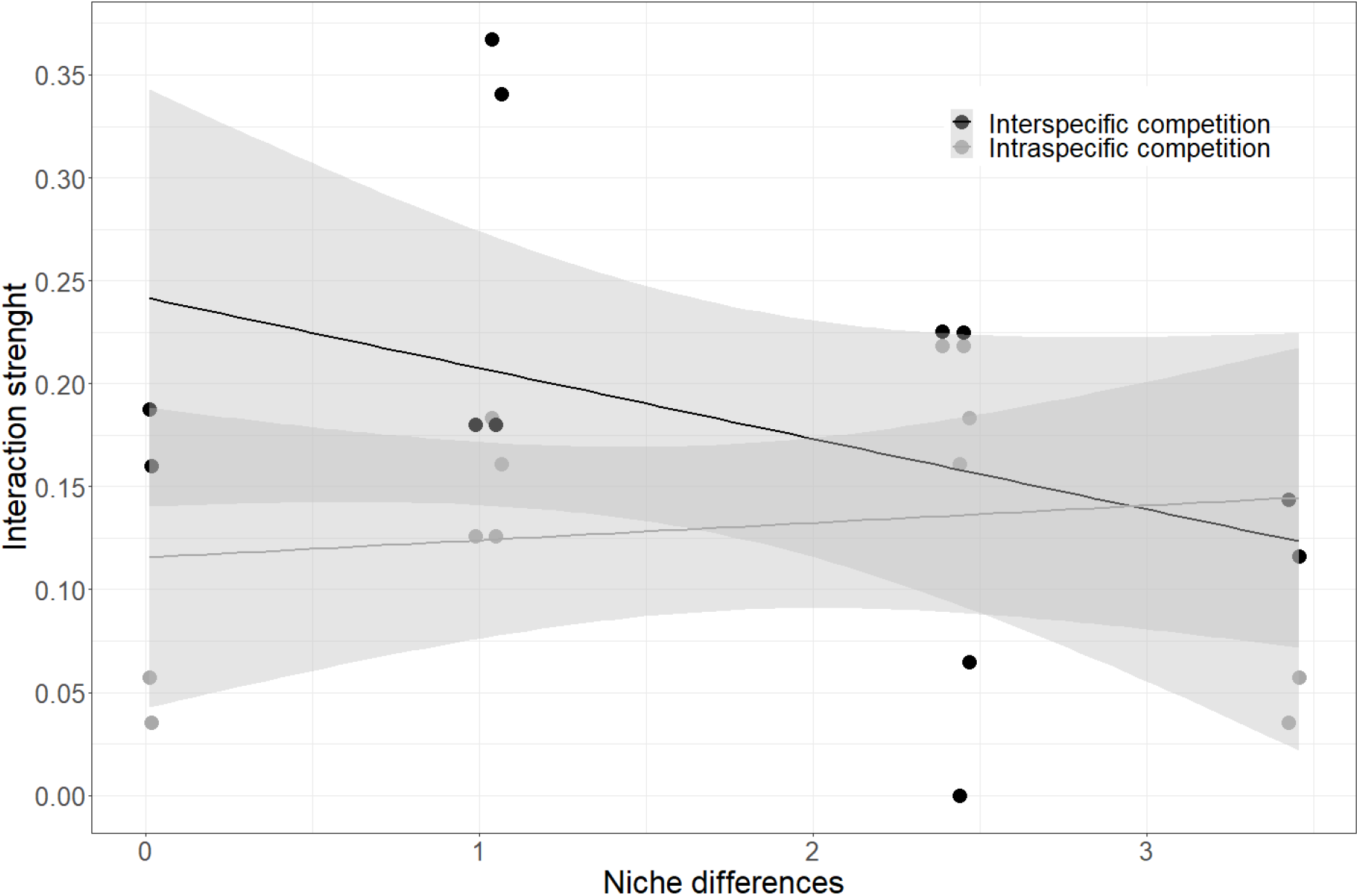
The effect of intra and interspecific interactions in driving variation in niche differences between genotype pairs.

We did not observe that differences between these two clusters were attributable to a particular genotypic difference. For both tetraploid and hexaploid groups, we observed pairs of genotypes that differed in the ploidy level as well as their degree of heterozygosity (e.g. see interaction between 4xOC and 6xS in priority effect group and 6xOC with 4xS in coexistence group) and others that did only differ in one aspect (e.g. see interaction between 4xOC and 4xS in coexistence group and 4xS and 6xS in priority effects group) (Fig. 3). Despite this variability, we found that a particular vegetative character allows differentiating these two groups of pairs of genotypes. Specifically, the number of stalks, which is a character related to the size of the plant, predicted differences observed in competitive outcomes between coexistence and priority effects (Multiple Analysis of Covariances F = 39.91, p < 0.01). Genotypes with larger numbers of stalks also showed priority effects (Fig 5), suggesting that space is a critical resource for which these genotypes compete.

**Figure 5.**
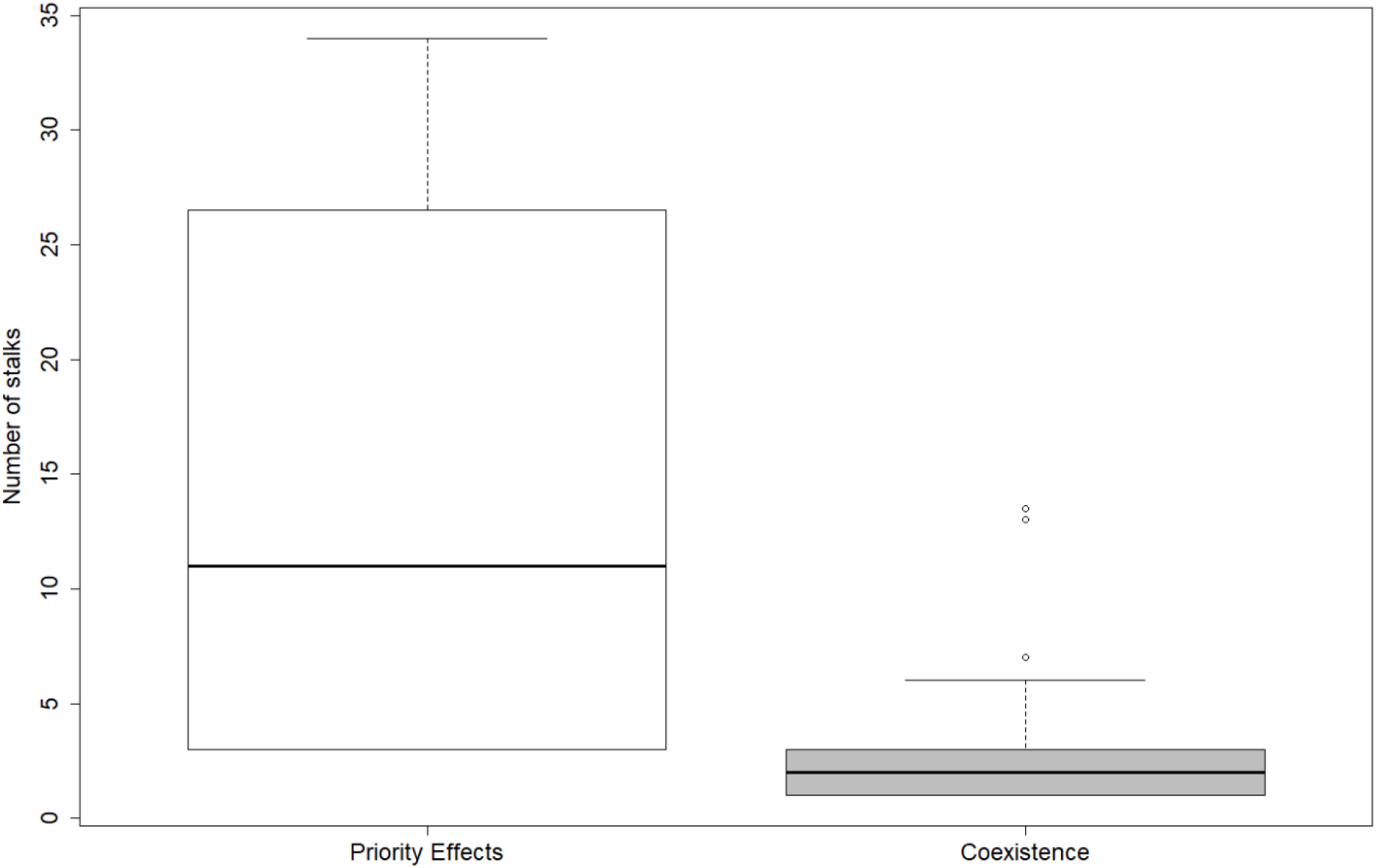
Boxplot represents median and quartiles of number of stalks for both coexistence outcomes we predicted when combining ploidy level and the degree of heterozygosity. Priority effects correspond to the combination of the three following genetic pairs: 4xS with 6xS, 4xOC with 6xOC and 4xOC with 6xS, while Coexistence groups the other three genetic pairs: 4xS with 6xOC, 4xOC with 4xS and 6xOC and 6xS.

## Discussion

Understanding the ecological consequences of genetic diversity is key to explain observed patterns of sympatric and allopatric genetic populations in nature. However, this understanding has been seldom explored because there is a lack of connection between ecological theory that describes the dynamics of interacting organisms and genetic material that can be manipulated to assess competitive interactions. In this study, we show how genetic diversity mechanisms provide a wide range of ecological outcomes based on the strength of competitive interactions between genotypes. Contrary to our expectations, low ploidy level, showed higher competitive ability, compared with tetraploid and hexaploid plants of *E. incanum*, mainly due to a lower response to competition and to a higher seed production to a lesser extent. That is, the intrinsic ability of diploids to produce seeds was little affected by increases in the density of individuals of the same and different genotypes within the community. Although theory commonly predicts that an increase in ploidy should confer an increase in fitness, these competitive advantages tend to occur under changing or extreme environments due to aridity or cold (López-Jurado et al. 2016; Liu et al. 2021). Instead, in our experiment, conditions were stable and non-stressful (i.e. no drought treatment). Under such conditions recent computational work simulating biological evolution suggested that non-polyploid perform better than polyploids (Yao *et al*., 2019; Carretero-Paulet & Van de Peer, 2020). This phenomenon can be explained by the amplification of the effect of random mutations on their gene regulatory networks because of the rise of complexity linked to whole genome duplication (multiplying the number of nodes and interaction in the gene regulatory network). Random mutation, often maladaptive or detrimental, under stable environments will propagate widely. In contrast, a stressful or unstable environment may provide substantive variation for survival (Yao et al. 2019; Carretero-Paulet and Van de Peer 2020; Van de Peer et al. 2021).

The reason for why polyploids were more sensitive to reduce their fitness in terms of seed production in the presence of neighbors can be due to the fact that they incur in a trade-off between being adapted to stressful conditions and tolerating competition from neighbors (López-Jurado et al. 2019; López-Jurado et al. 2016). In fact, it has been found under natural conditions that polyploids adapted to arid conditions present in general low densities (Manzaneda *et al*., 2012; Penner *et al*., 2020). Nevertheless, the higher competitive ability we observed in non-polyploids was not translated to predict competitive exclusion between diploids and the rest of genotypes (tetraploids and hexaploids). The structure of intra and interspecific interactions between genetic pairs modulate this expectation to produce strong priority effects instead, meaning that the historical contingency such as the genotype that arrives first to the community excludes the other.

Our results, therefore, suggest that certain evolutionary theories conveying a competitive advantage to more diverse genotypes should be revisited. In that sense, current theory puts a major role in the effect of environments on driving an increase of ploidy and associated competitive advantages and the ability to colonize and dominate novel stressful environments. However, these increases in ploidy does not always occur as exemplifies diploids of *Erysimum mediohispanicum* (another species belonging to *Erysimum*), which have been associated with hard conditions at high altitudes (Muñoz-Pajares et al., 2018). Even if they occur, such advantages seem to be related to species vital rates such as seed production or survival but it comes at the trade-off to tolerate heterospecific competition. This lack of ability to tolerate competition might compromise the successful colonization of new habitats to contingency events as we observed in the priority effects case, or it might be also the case of being excluded from communities that are well established with high local abundances. Further experiments across environmental gradients are needed to reveal the consequences of increases in ploidy for determining trade-off between seed production and tolerance to competition, and therefore, to tease apart the role of the abiotic component (stress conditions) from the biotic component (competition) in driving polyploids advantages to colonize novel environments.

Although most of the work on evolutionary biology has studied the role of variation in ploidy in promoting differences in fitness between species, much less has been explored its effect in promoting niche differences between genotypes (Alonso-Marcos et al. 2019; Hülber et al. 2018; Balao et al. 2011). Our results strongly suggest that ploidy variation promotes the demographic consequences of niche differences which stabilize the population dynamics of competing genotypes and further indicate that this effect is in turn strongly influenced by the degree of heterozygosity. Specifically, we found that both levels of heterozygosity can coexist within the same level of ploidy (this is true for tetraploids and hexaploids) as well as between levels of ploidy. This result, which goes against predictions that heterozygote should exclude homozygous genotypes as well as higher ploidy should exclude lower ploidy genotypes, is very important because it suggests that both sources, ploidy and heterozygosity, are critical to maintain genetic diversity within and across genotypes. Moreover, with our ability to link the strength of genotype interactions with their likelihood to coexist, we found that genotypes were not weakly differentiated, as we might expect from closely related evolutionary units within the *E. incanum* system. That is, in those genetic pairs predicted to coexist, we did not observe that weak niche differences overcome small fitness differences. Instead, they presented strong niche differentiation. Such high niche differences indicate that genotypes experience greater intraspecific than interspecific competition. Although with our experiment it is not possible to know the ultimate sources of these axes promoting niche differences, we were not expecting these results considering that the greenhouse experiment was settled in relatively small pots where there were few environmental axes compared with natural conditions.

Besides coexistence, we also predicted that several of these tetra and hexaploids genotypes would incur in priority effects according to our experiment conditions. These priority effects mean that genotypes experienced positive rather than negative density dependence. That is, genotypes favor themselves rather than limit their competitors and the genotype exclusion is not due to deterministic processes but rather due to historical contingency such as order of arrival. At a single location, this contingency promotes the dominance of the genotype that arrives first, but at larger scales coexistence can occur if both genotypes arrive first to different locations. If we take a closer look at these genotypes incurring in priority effects, we observed that tetraploids with high levels of heterozygosity (4xOC see Fig. 3) are able to exclude any hexaploid plant if they arrive first. This result is very interesting as these species belong to a selfing clade (Abdelaziz *et al*., 2019) and the mating system transition theories predict that once selfing populations have purged its genetic load, no advantage of high heterozygous plants is expected (Goodwillie *et al*., 2005). However, the overdominance genetic model predicts that heterozygotes would be superior to homozygotes at loci affecting fitness (Khotyleva et al., 2017).

Based on independent information of either the level of ploidy or the degree of heterozygosity, we could not differentiate those genetic pairs predicted to coexist from those incurring in priority effects. However, we found that the phenotypic expression of these genotypes, measured as the number of stalks, differentiated these contrasted coexistence outcomes. Presenting a high number of stalks is an important feature that equally favors competitive ability by promoting low response to competition, and high plant performance in terms of seed production. Therefore, genotypes with greater number of stalks can incur in priority effects for the following reasons. On the one hand, the production of more flowering stalks or developing them faster allows genotypes to occupy more space, and compete better by monopolizing more resources and by shadowing other surrounding neighbors (Craine & Dybzinski, 2013). On the other hand, plants, as modular organisms, the bigger they are, the more reproductive organs they develop. Thus, the number of flowering stalks has a direct effect on plant fitness by its effect on the number of flowers produced. This is the case of different species which use the size of plants to attract pollen vectors (Klinkhamer *et al*., 1989; Klinkhamer & de Jong, 1990)), including species in the *Erysimum* genus (Gómez *et al*., 2009; Alonso-Marcos *et al*., 2019)). But the production of more flowers also means higher fitness values when the plant has the ability to self-pollinate (Gerber, 1985). However, why genotypes with high number of stalks incur in positive density dependence (priority effects) and why those with low number of stalks do the opposite (coexistence) is unclear. Unfortunately, there is no prior study in the literature that has found a single trait driving the strong variation in niche differences from negative to positive as we found in our experiment. It might be the case that the number of stalks correlate with other traits promoting niche differences such as differences in phenology (Navas and Violle 2009; Godoy and Levine 2014), of the ability to make photosynthesis at different light irradiances (Pérez-Ramos *et al*., 2019). Further studies need to explore more in depth the multiple and correlated phenotypic expression and associated mechanisms that underlie these switches between positive and negative density-dependence processes, but our results are the first to highlight that variation in the number of stalks, as a subrogate of plant size, is critical to predict opposed coexistence outcomes by varying niche differences among taxa.

Our study provides strong evidence that genetic diversity plays a critical role in determining ecological outcomes between closely related genotypes. Contrary to expectations, diploids showed greater competitive ability than tetra and hexaploids. However, this competitive advantage did not translate to competitive dominance and the exclusion of the inferior competitors, rather, we predicted that the winner of competition depends on contingency such as the genotype that arrives first to a location. This is an interesting result that needs further consideration in future work because priority effects are not an ecological outcome considered by current theories in evolution describing the advantage of polyploids and the consequences of polyploidization for describing allopatric and sympatric populations. Moreover, we found that ploidy interacts with the degree of heterozygosity to reverse the competitive outcomes from priority effects to coexistence, which highlights the importance of keeping a diverse genetic background within genotypes to maintain in turn genetic diversity across genotypes. Although linking genetic diversity with competition outcomes can be difficult for logistical and methodological limitations, our results strongly suggest that the phenotypic expression of an easy-to-measure trait, the number of stalks, can predict such variation in ecological outcomes. In particular, pairs of genotypes that show on average more stalks incur in priority effects, whereas those with low stalks number are predicted to coexist. This study does not explore why the number of stalks, considered a subrogate of competition for space, can change competition from positive to negative density dependence (negative versus positive niche differences). Yet, our results highlight the importance of exploring the effects of genetic diversity on the interactions among genotypes because they can strongly modify their ecological dynamics compared to expectations from only responses to the environment.

## Acknowledgements

The authors thank to Modesto Berbel, Celia Vaca-Benito, Andrea Martín-Salas, María de la Paz Solís for their help to set up and conduct the greenhouse experiment. It was not an easy task during the COVID-19 pandemic. This work was funded by grants *OUTevolution* from the Spanish Ministry of Science and Innovation (PID2019-111294GB-I00/SRA/10.13039/501100011033), *globalHybrids* from the *Organismo Autónomo de Parques Nacionales* (Ref: 2415/2017). AG-M was supported by *OUTevolution* project (PID2019-111294GB-I00/SRA/10.13039/501100011033). OG acknowledges the postdoctoral financial support provided by the Spanish Ministry of Economy and Competitiveness (MINECO) and by the European Social Fund through the Ramon y Cajal Programme (RYC-2017-23666). J.A. Pérez-Romero thanks the Ministerio de Ciencia y Educación for his personal financial support (FJC2020-043865-I).

## Authors contribution

O.G and M.A conceptualized the experiment. A.G-M and M.A contributed to conduct the experiment and data collection. J.A.P-R led modeling and statistical analyses. J.A.P-R and O.G wrote the first draft of the manuscript and all authors provided substantial revisions to the manuscript.

## Data availability

Data and code is storage in https://drive.google.com/drive/folders/1dCU49U7GOHEd3xdO6xGeUFND-Tki5pZq?usp=share_link for review purposes, and will make it available in a public repository upon acceptance.

## References

Abdelaziz M, Bakkali M, Gómez JM, Olivieri E, Perfectti F. 2019. Anther Rubbing, a New Mechanism That Actively Promotes Selfing in Plants. The American naturalist 193: 140–147.

Abdelaziz M, Lorite J, Muñoz-Pajares AJ, Herrador MB, Perfectti F, Gómez JM. 2011. Using complementary techniques to distinguish cryptic species: a new Erysimum (Brassicaceae) species from North Africa. American journal of botany 98: 1049–1060.

Adler PB, Hillerislambers J, Levine JM. 2007. A niche for neutrality. Ecology letters 10: 95–104.

Alonso-Marcos H, Nardi FD, Scheffknecht S, Tribsch A, Hülber K, Dobeš C. 2019. Difference in reproductive mode rather than ploidy explains niche differentiation in sympatric sexual and apomictic populations of. Ecology and evolution 9: 3588–3598.

Al-Shehbaz IA, Beilstein MA, Kellogg EA. 2006. Systematics and phylogeny of the Brassicaceae (Cruciferae): an overview. Plant systematics and evolution = Entwicklungsgeschichte und Systematik der Pflanzen 259: 89–120.

Ayala FJ, Campbell CA. 1974. Frequency-Dependent Selection. Annual Review of Ecology and Systematics 5: 115–138.

Balao F, Herrera J, Talavera S. 2011. Phenotypic consequences of polyploidy and genome size at the microevolutionary scale: a multivariate morphological approach. The New phytologist 192: 256–265.

Bartomeus I, Saavedra S, Rohr RP, Godoy O. 2021. Experimental evidence of the importance of multitrophic structure for species persistence. Proceedings of the National Academy of Sciences of the United States of America 118.

te Beest M, Le Roux JJ, Richardson DM, Brysting AK, Suda J, Kubesová M, Pysek P. 2012. The more the better? The role of polyploidy in facilitating plant invasions. Annals of botany 109: 19–45.

Bichet C, Vedder O, Sauer-Gürth H, Becker PH, Wink M, Bouwhuis S. 2019. Contrasting heterozygosity-fitness correlations across life in a long-lived seabird. Molecular ecology 28: 671–685.

Bird KA, VanBuren R, Puzey JR, Edger PP. 2018. The causes and consequences of subgenome dominance in hybrids and recent polyploids. The New phytologist 220: 87–93.

Bomblies K, Weigel D. 2007. Hybrid necrosis: autoimmunity as a potential gene-flow barrier in plant species. Nature reviews. Genetics 8: 382–393.

Carretero-Paulet L, Van de Peer Y. 2020. The evolutionary conundrum of whole-genome duplication. American journal of botany 107: 1101–1105.

Charlesworth B, Charlesworth D. 1999. The genetic basis of inbreeding depression. Genetical Research 74: 329–340.

Chen ZJ, Ni Z. 2006. Mechanisms of genomic rearrangements and gene expression changes in plant polyploids. BioEssays: news and reviews in molecular, cellular and developmental biology 28: 240–252.

Chesson P. 2000. Mechanisms of maintenance of species diversity. Annual review of ecology and systematics 31: 343–366.

Chesson P. 2012. Species Competition and Predation. Encyclopedia of Sustainability Science and Technology: 10061–10085.

Comai L. 2005. The advantages and disadvantages of being polyploid. Nature reviews. Genetics 6: 836–846.

Conover JL, Wendel JF. 2022. Deleterious Mutations Accumulate Faster in Allopolyploid Than Diploid Cotton (Gossypium) and Unequally between Subgenomes. Molecular biology and evolution 39.

DeMalach N, Zaady E, Kadmon R. 2017. Light asymmetry explains the effect of nutrient enrichment on grassland diversity. Ecology letters 20: 60–69.

Dempster ER. 1955. MAINTENANCE OF GENETIC HETEROGENEITY. Cold Spring Harbor Symposia on Quantitative Biology 20: 25–32.

Ellegren H, Sheldon BC. 2008. Genetic basis of fitness differences in natural populations. Nature 452: 169–175.

Fennane M, Ibn-Tattou M. 1999. Flore pratique du Maroc. Vol. 1. Pteridophyta, Gymnospermae, Angiospermae (Lauraceae-Neuradaceae). Inst. Scientifique.

Fragata I, Costa-Pereira R, Kozak M, Majer A, Godoy O, Magalhães S. 2022. Specific sequence of arrival promotes coexistence via spatial niche pre-emption by the weak competitor. Ecology letters 25: 1629–1639.

Fridman E. 2015. Consequences of hybridization and heterozygosity on plant vigor and phenotypic stability. Plant science: an international journal of experimental plant biology 232: 35–40.

García-Callejas D, Godoy O, Bartomeus I. 2020. cxr : A toolbox for modelling species coexistence in R. Methods in Ecology and Evolution 11: 1221–1226.

García-Muñoz A, Ferrón C, Vaca-Benito C, Loureiro J, Castro S, Jesús Muñoz-Pajares A, Abdelaziz M. Ploidy effects on the relationship between floral phenotype, reproductive investment and fitness exhibited by an autogamous species complex.

Gerber MA. 1985. The Relationship of Plant Size to Self-Pollination in Mertensia Ciliata. Ecology 66: 762–772.

Germain RM, Weir JT, Gilbert B. 2016. Species coexistence: macroevolutionary relationships and the contingency of historical interactions. Proceedings. Biological sciences / The Royal Society 283: 20160047.

Gibson DJ, Connolly J, Hartnett DC, Weidenhamer JD. 1999. Designs for greenhouse studies of interactions between plants. Journal of Ecology 87: 1–16.

Godoy O, Kraft NJB, Levine JM. 2014. Phylogenetic relatedness and the determinants of competitive outcomes. Ecology letters 17: 836–844.

Godoy O, Levine JM. 2014. Phenology effects on invasion success: insights from coupling field experiments to coexistence theory. Ecology 95: 726–736.

Gómez JM, Perfectti F, Bosch J, Camacho JPM. 2009. A geographic selection mosaic in a generalized plant–pollinator–herbivore system. Ecological Monographs 79: 245–263.

Goodwillie C, Kalisz S, Eckert CG. 2005. The Evolutionary Enigma of Mixed Mating Systems in Plants: Occurrence, Theoretical Explanations, and Empirical Evidence. Annual review of ecology, evolution, and systematics 36: 47–79.

Grant V. 1981. Plant Speciation. Columbia University Press.

Harpole WS, Sullivan LL, Lind EM, Firn J, Adler PB, Borer ET, Chase J, Fay PA, Hautier Y, Hillebrand H, et al. 2016. Addition of multiple limiting resources reduces grassland diversity. Nature 537: 93–96.

Hart SP, Freckleton RP, Levine JM. 2018. How to quantify competitive ability. Journal of Ecology 106: 1902–1909.

Hart SP, Turcotte MM, Levine JM. 2019. Effects of rapid evolution on species coexistence. Proceedings of the National Academy of Sciences of the United States of America 116: 2112–2117.

Hayes HK, Others. 1952. Development of the heterosis concept. Heterosis: 49–65.

Hernández-Leal MS, Suárez-Atilano M, Piñero D, González-Rodríguez A. 2019. Regional patterns of genetic structure and environmental differentiation in willow populations (Salix humboldtiana Willd.) from Central Mexico. Ecology and evolution 9: 9564–9579.

Hughes AR, Inouye BD, Johnson MTJ, Underwood N, Vellend M. 2008. Ecological consequences of genetic diversity. Ecology letters 11: 609–623.

Hülber K, Sonnleitner M, Haider J, Schwentenwein M, Winkler M, Schneeweiss GM, Schönswetter P. 2018. Reciprocal transplantations reveal strong niche differentiation among ploidy-differentiated species of the Senecio carniolicus aggregate (Asteraceae) in the easternmost Alps. Alpine Botany 128: 107–119.

Jürgens A, Witt T, Gottsberger G. 2002. Pollen grain numbers, ovule numbers and pollen-ovule ratios in Caryophylloideae: correlation with breeding system, pollination, life form, style number, and sexual system. Sexual plant reproduction 14: 279–289.

Ke P-J, Letten AD. 2018. Coexistence theory and the frequency-dependence of priority effects. Nature ecology & evolution 2: 1691–1695.

Klinkhamer PGL, de Jong TJ. 1990. Effects of Plant Size, Plant Density and Sex Differential Nectar Reward on Pollinator Visitation in the Protandrous Echium vulgare (Boraginaceae). Oikos 57: 399.

Klinkhamer PGL, de Jong TJ, de Bruyn G-J. 1989. Plant Size and Pollinator Visitation in Cynoglossum Officinale. Oikos 54: 201.

Kolář F, Lučanová M, Vít P, Urfus T, Chrtek J. 2013. Diversity and endemism in deglaciated areas: ploidy, relative genome size and niche differentiation in the Galium pusillum complex (Rubiaceae) in Northern and …. Annals of.

Lankau RA, Nuzzo V, Spyreas G, Davis AS. 2009. Evolutionary limits ameliorate the negative impact of an invasive plant. Proceedings of the National Academy of Sciences of the United States of America 106: 15362–15367.

Lavorel S, Garnier E. 2002. Predicting changes in community composition and ecosystem functioning from plant traits: revisiting the Holy Grail. Functional ecology 16: 545–556.

Leebens-Mack JH, Barker MS, Carpenter EJ, Deyholos MK, Gitzendanner MA, Graham SW, Grosse I, Li Z, Melkonian M, Mirarab S, et al. 2019. One thousand plant transcriptomes and the phylogenomics of green plants. Nature 574: 679–685

Levin DA. 1981. Dispersal Versus Gene Flow in Plants. Annals of the Missouri Botanical Garden. Missouri Botanical Garden 68: 233–253.

Levine JM, HilleRisLambers J. 2009. The importance of niches for the maintenance of species diversity. Nature 461: 254–257.

Levin DA, Levin of IBD. 2002. The Role of Chromosomal Change in Plant Evolution. Oxford University Press.

Liston A, Wei N, Tennessen JA, Li J, Dong M, Ashman T-L. 2020. Revisiting the origin of octoploid strawberry. Nature genetics 52: 2–4.

Liu J, Li J, Fu C. 2021. Comparative physiology and transcriptome analysis reveals the regulatory mechanism of genome duplication enhancing cold resistance in Fragaria nilgerrensis. Environmental and Experimental Botany 188: 104509.

López-Jurado J, Balao F, Mateos-Naranjo E. 2016. Deciphering the ecophysiological traits involved during water stress acclimation and recovery of the threatened wild carnation, Dianthus inoxianus. Plant physiology and biochemistry: PPB / Societe francaise de physiologie vegetale 109: 397–405.

López-Jurado J, Mateos-Naranjo E, Balao F. 2019. Niche divergence and limits to expansion in the high polyploid Dianthus broteri complex. The New phytologist 222: 1076–1087.

Manzaneda AJ, Rey PJ, Bastida JM, Weiss-Lehman C, Raskin E, Mitchell-Olds T. 2012. Environmental aridity is associated with cytotype segregation and polyploidy occurrence in Brachypodium distachyon (Poaceae). The New phytologist 193: 797–805.

Meudt HM, Albach DC, Tanentzap AJ, Igea J, Newmarch SC, Brandt AJ, Lee WG, Tate JA. 2021. Polyploidy on Islands: Its Emergence and Importance for Diversification. Frontiers in plant science 12: 637214.

Moghe GD, Shiu S-H. 2014. The causes and molecular consequences of polyploidy in flowering plants. Annals of the New York Academy of Sciences 1320: 16–34.

Narwani A, Alexandrou MA, Oakley TH, Carroll IT, Cardinale BJ. 2013. Experimental evidence that evolutionary relatedness does not affect the ecological mechanisms of coexistence in freshwater green algae. Ecology letters 16: 1373–1381.

Navas ML, Violle C. 2009. Plant traits related to competition: how do they shape the functional diversity of communities? Community Ecology 10: 131–137.

Nieto Feliner G. 2014. Patterns and processes in plant phylogeography in the Mediterranean Basin. A review. Perspectives in plant ecology, evolution and systematics 16: 265–278.

Nieto Feliner G, Casacuberta J, Wendel JF. 2020. Genomics of Evolutionary Novelty in Hybrids and Polyploids. Frontiers in genetics 11: 792.

Otto SP, Whitton J. 2000. Polyploid incidence and evolution. Annual review of genetics 34: 401–437.

Parisod C, Holderegger R, Brochmann C. 2010. Evolutionary consequences of autopolyploidy. The New phytologist 186: 5–17.

Paterson AH. 2005. Polyploidy, evolutionary opportunity, and crop adaptation. Genetica 123: 191–196.

Penner S, Dror B, Aviezer I, Bar-Lev Y, Salman-Minkov A, Mandakova T, Šmarda P, Mayrose I, Sapir Y. 2020. Phenology and polyploidy in annual Brachypodium species (Poaceae) along the aridity gradient in Israel. Journal of systematics and evolution 58: 189–199.

Pérez-Ramos IM, Matías L, Gómez-Aparicio L, Godoy ó. 2019. Functional traits and phenotypic plasticity modulate species coexistence across contrasting climatic conditions. Nature communications 10: 2555.

Petry WK, Kandlikar GS, Kraft NJB, Godoy O, Levine JM. 2018. A competition–defence trade-off both promotes and weakens coexistence in an annual plant community. Journal of Ecology 106: 1806–1818.

R Core Team, R., & R Core Team. 2021. R: a language and environment for statistical computing. R Foundation for Statistical Computing; 2020.

Raabová J, Fischer M, Münzbergová Z. 2008. Niche differentiation between diploid and hexaploid Aster amellus. Oecologia 158: 463–472.

Ramsey J, Ramsey TS. 2014. Ecological studies of polyploidy in the 100 years following its discovery. Philosophical transactions of the Royal Society of London. Series B, Biological sciences 369.

Ramsey J, Schemske DW. 2002. Neopolyploidy in Flowering Plants. Annual Review of Ecology and Systematics 33: 589–639.

Rey PJ, Manzaneda AJ, Alcántara JM. 2017. The interplay between aridity and competition determines colonization ability, exclusion and ecological segregation in the heteroploid Brachypodium distachyon species complex. The New phytologist 215: 85–96.

Salmon A, Ainouche ML, Wendel JF. 2005. Genetic and epigenetic consequences of recent hybridization and polyploidy in Spartina (Poaceae). Molecular ecology 14: 1163–1175.

Soltis DE, Albert VA, Leebens-Mack J, Bell CD, Paterson AH, Zheng C, Sankoff D, Depamphilis CW, Wall PK, Soltis PS. 2009. Polyploidy and angiosperm diversification. American journal of botany 96: 336–348.

Soltis DE, Soltis PS. 1999. Polyploidy: recurrent formation and genome evolution. Trends in ecology & evolution 14: 348–352.

Sonnleitner M, Flatscher R, Escobar García P, Rauchová J, Suda J, Schneeweiss GM, Hülber K, Schönswetter P. 2010. Distribution and habitat segregation on different spatial scales among diploid, tetraploid and hexaploid cytotypes of Senecio carniolicus (Asteraceae) in the Eastern Alps. Annals of botany 106: 967–977.

Tilman C, Sandhu R. 1998. A model recycling program for Alabama. Resources, Conservation and Recycling 24: 183–190.

Van de Peer Y, Ashman T-L, Soltis PS, Soltis DE. 2021. Polyploidy: an evolutionary and ecological force in stressful times. The Plant cell 33: 11–26.

Violle C, Garnier E, Lecoeur J, Roumet C, Podeur C, Blanchard A, Navas M-L. 2009. Competition, traits and resource depletion in plant communities. Oecologia 160: 747–755.

Wang X, Shi X, Hao B, Ge S, Luo J. 2005. Duplication and DNA segmental loss in the rice genome: implications for diploidization. The New phytologist 165: 937–946.

Wickham H. 2016. ggplot2: Elegant Graphics for Data Analysis. Springer.

Wilke, C. O. 2019. cowplot: Streamlined Plot Theme and Plot Annotations for ‘ggplot2’; 2020. R package version, 1(1).

Wood TE, Takebayashi N, Barker MS, Mayrose I, Greenspoon PB, Rieseberg LH. 2009. The frequency of polyploid speciation in vascular plants. Proceedings of the National Academy of Sciences of the United States of America 106: 13875–13879.

Yaakov B, Kashkush K. 2011. Methylation, transcription, and rearrangements of transposable elements in synthetic allopolyploids. International journal of plant genomics 2011: 569826.

Yao Y, Carretero-Paulet L, Van de Peer Y. 2019. Using digital organisms to study the evolutionary consequences of whole genome duplication and polyploidy. PloS one 14: e0220257.

